# Cooperation Between Genotoxic Bacteria Accelerates Tumorigenesis in a Mouse Model of Colon Carcinogenesis

**DOI:** 10.1101/2025.04.18.649375

**Authors:** Tamar Plitt, Alix Piessevaux, Urvija Rajpal, Jeremy Fisher, Matthew P. Spindler, Zhihua Li, Ilaria Mogno, Ye Yang, A. Nicole Desch, Gerald Chu, Zhengyu Jiang, Jianyao Wang, Dirk Gevers, David Pocalyko, Abby L. Geis, Christian Jobin, Kurtis E. Bachman, Cynthia L. Sears, Graham J. Britton, Lani R. San Mateo, Jeremiah J. Faith

## Abstract

To identify causal links between gut microbes and tumorigenesis, we colonized groups of germ-free colon tumor-susceptible mice (*Apc*^*Min/+*^;*Il10*^−/−^) with 15 cultured human fecal microbiotas from healthy individuals, as well as patients with inflammatory bowel disease and colorectal cancer. The number of colonic tumors in *Apc*^*Min/+*^;*Il10*^−/−^ mice varied by donor microbiota but not by the health status of the donor. In vitro screens of host cell proliferation, genotoxicity, and inflammation in bacteria-mammalian cell cocultures revealed that genotoxicity best predicted tumorigenic microbes in vivo with genotoxic microbes present in all tested individuals. The genotoxic subset of strains from each donor induced more tumors than the complete community – even when the complete community was not tumorigenic. Combining genotoxic microbes from multiple sources increased tumor number and decreased the time to tumor onset. Together these results suggest that most individuals harbor genotoxic bacterial strains and the balance of genotoxic to protective strains determines the timing and severity of tumorigenesis in vivo.

## INTRODUCTION

Colorectal cancer (CRC) is the second leading cause of cancer-related deaths worldwide, with rising incidence in individuals less than 50 years old^1–3^. As with many diseases, both genetic and environmental factors contribute to disease development. However, genetic predisposition for CRC only accounts for 2-5% of all CRC cases, highlighting the much larger role of the environment in driving sporadic CRC^4,5^.

Perturbations in the composition and metabolic output of the gut microbiota are associated with diseases ranging from immune-mediated diseases to metabolic diseases to cancer. Given that the colon is one of most densely populated microbial ecosystems on our planet, the contribution of microorganisms to colorectal carcinogenesis has been increasingly recognized^6–8^. Metagenomic studies in humans have shown significant differences in the gut microbiota composition of CRC patients compared to the healthy population, in terms of species richness and the abundance of potentially protective and/or pro-carcinogenic taxa^2,9–15^. In addition, several germ-free (GF) mouse models of CRC are devoid of tumors until after they are colonized by complex microbiota from human or mouse stool^9,10,16–18^. Human colon mucosal microbes have also been associated with CRC pathogenesis, as colon biofilms from CRC patients or healthy individuals undergoing routine colonoscopies revealed tumorigenic microbes in murine models of disease that retained tumorigenic capacity upon transfer to additional GF mice^18^.

Emerging evidence suggests diverse mechanisms by which the gut microbiota may lead to CRC^8,19–27^. Examples include CRC-associated *Fusobacterium nucleatum* strains that enhance tumor growth by activating pro-inflammatory NF-kB signaling pathway within the tumor microenvironment through pathogen recognition receptor (PRR) engagement^28–30^ or via pro-proliferative signaling pathways through FadA binding of E-cadherin on epithelial cells^31^. Another CRC-associated human commensal bacterium, enterotoxigenic *Bacteroides fragilis* (ETBF), triggers colitis and colonic tumors in a Stat3- and Th17-dependent manner in CRC-susceptible mice^32^. Beyond proliferation and inflammation, *Escherichia coli* expressing the genomic island polyketide synthase (*pks*^+^) produce the small-molecule genotoxin colibactin that alkylates adenine residues causing DNA damage within the epithelial cells leading to chromosomal instability^33,34^. There is great interest in studying the role of *pks*^+^ *E. coli* in the colorectal carcinogenesis as these strains have been detected in human CRCs and possess the ability to potentiate tumorigenesis in animal models of disease^17,35,36^. ETBF has also been shown to damage the host DNA through induction of reactive oxygen species (ROS)^37,38^. Recently, indolimine-producing *M. morganii* was discovered to induce genotoxicity and exacerbate tumorigenesis in gnotobiotic mice through increased intestinal permeability^39^. The interplay between the gut microbiome and cancer remains only partially characterized, in part due to the lack of high-throughput scalable platforms to identify the functions of the understudied majority of strains in the microbiome^40^.

In this study, we use a CRC-susceptible gnotobiotic mouse model colonized with human microbiota to identify causal links between gut microbes, colitis susceptibility, and tumorigenesis. We evaluate the tumorigenic potential of individual human gut microbiota from CRC, inflammatory bowel disease (IBD), and healthy donors (HDs) in ex-GF *Apc*^*Min/+*^*;Il10*^−/−^ mice and find large interpersonal variation on colonic tumor burden across the general population, regardless of the health status of the originating donor. In parallel, we use scalable high-throughput in vitro platforms to characterize the function of hundreds of microbial strains across the general population to explore the microbiome’s influence on colon carcinogenesis by promoting inflammation, proliferation, or genotoxicity within intestinal epithelial cells. We find a broad range of strain-specific induction of inflammation, proliferation, and genotoxicity across tumorigenic and non-tumorigenic donors. However, only the genotoxic microbial subsets predict distal colon tumorigenesis. Finally, we show that combinations of pathogenic bacteria, in the absence of protective bacteria, exacerbate tumor burden and accelerate tumor onset. Together, these data reveal new insights into the causal relationships between the gut microbiome and cancer and characterize the tumorigenesis-related functions of hundreds of strains in the microbiome.

## RESULTS

### Human gut microbiotas differentially drive tumorigenesis in susceptible mice

To investigate the tumorigenic potential of human gut microbiotas, we colonized 15 cultured human gut microbiota isolated from stool samples of CRC (N=4), IBD (N=4; Crohn’s disease=2, ulcerative colitis=2), and HDs (N=7) in separate groups of GF *Apc*^*Min/+*^*;Il10*^−/−^ mice, notable for their microbiota-dependent susceptibility to spontaneous tumors in the colon (Figure 1A)^10^. We used cultured, defined polymicrobial communities to precisely control the composition of the microbiotas and to enable dissection of specific causative strains in vitro and in vivo. We found that different human cultured gut microbiotas induced significantly different number of tumors (p<0.001, Kruskal-Wallis; Figure 1B) regardless of the donor’s health status (p=0.906, Kruskal-Wallis; Figure S1A), with several microbiotas leading to no tumors at all (Figure 1B). CRC1, IBD3, IBD4, HD2, HD3, and HD5 microbiotas led to a similar number of tumors as those found in positive control mice mono-colonized with known tumorigenic bacterial strains ETBF (piglet 86-5443-2-2) or *pks*^+^ *E. coli* (murine strain NC101)^17,32,37^. Together, these results demonstrate the important role of interpersonal variation of human gut microbiota on colonic tumor burden.

**Fig. 1:**
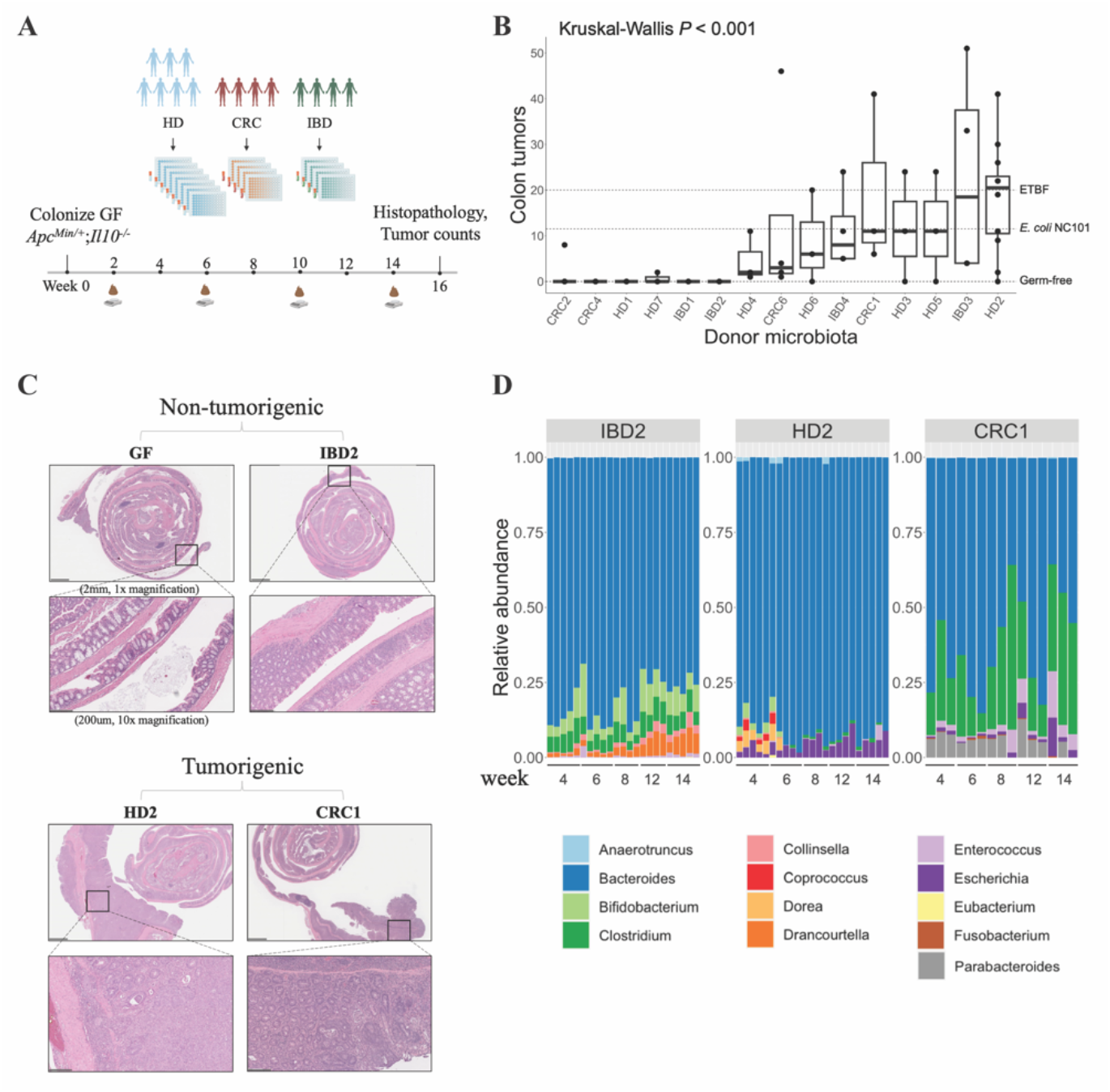
Human gut microbiotas differentially drive tumorigenesis in susceptible mice. **(A)** Overview of experimental outline. Figure created with BioRender.com. HD=Healthy donor, IBD=Inflammatory bowel disease, CRC=Colorectal cancer. **(B)** Macroscopic colon tumor counts from 20-22 week old Apc^Min/+^;Il10^−/−^ mice colonized with cultured microbiota collections from stools of CRC, IBD, and HDs. Each point represents data from one mouse. In the box plots: center line, median; box, IQR; whiskers, 1.5 × IQR. Statistical significance was calculated using Kruskal–Wallis. **(C)** Representative histological colon sections from mice colonized with non-tumorigenic (GF, IBD2) or tumorigenic (HD2, CRC1) microbiotas (5x, 40x magnification). Bottom panels represent insets of top panel boxed areas demonstrating normal intestinal tissue or development of intestinal neoplasia and inflammation. Scale bars = 2mm, 200μm. **(D)** Relative abundances of 13 genera by metagenomic profiling in mice colonized with the IBD2, HD2, and CRC1 microbiota across various time points (weeks post-gavage). The bars are colored according to genus name. N=3-5 mice per group.

Given the role of inflammation in the development and progression in CRC, we examined the histologic appearance of colons in mice colonized with cultured microbiota that led to tumor formations in vivo at the 16-week endpoint (termed “tumorigenic microbiota”) compared to microbiota that did not lead to tumor formations (termed “non-tumorigenic microbiotas”). Compared to GF mice or mice colonized with non-tumorigenic microbiotas (e.g., IBD2), *Apc*^*Min/+*^;*Il10*^−/−^ mice colonized with tumorigenic microbiotas (e.g., HD2, CRC1) developed moderate colitis with colonic neoplastic lesions, increased mononuclear inflammation in the lamina propria and submucosa, loss of goblet cells, crypt hyperplasia, and muscle thickening (Figure 1C).

In addition to terminal histological assessment of inflammation, we used fecal lipocalin-2 (LCN2)^41^ as a longitudinal measure of colitis severity. We found elevated fecal LCN2 in mice colonized with tumorigenic microbiotas compared to mice colonized with non-tumorigenic microbiotas (p=0.004, Mann-Whitney; Figure S1B). Tumor formation was significantly correlated with fecal LCN2 as early as 8 weeks post-colonization (p=0.004; Figure S1C).

Although these data support a potential role for inflammation in microbiota-specific tumor numbers, a number of mice with similar fecal LCN2 concentrations at 8 weeks post-colonization developed different numbers of tumors. These observations suggest that inflammation is not the sole contributing factor to tumorigenesis.

To understand if the observed inflammation was due to the colonization of colitis susceptible *Il10*^−/−^ mice with an inflammatory microbiome, we engrafted the tumorigenic HD2 microbiota into *Il10*^−/−^ mice containing the wild-type *Apc* allele with the non-tumorigenic IBD1 microbiota as a control. Colitis severity was monitored, and mice were sacrificed at 8 weeks post-colonization to assess intestinal inflammation. Intestinal inflammation, as measured by elevated fecal LCN2, was largely absent in *Il10*^−/−^ mice colonized with the tumorigenic HD2 microbiota compared to HD2-colonized *Apc*^*Min/+*^;*Il10*^−/−^ mice (p=0.016; Figure S1D), suggesting tumorigenesis is required to observe the increased intestinal inflammation in *Apc*^*Min/+*^;*Il10*^−/−^ mice.

Metagenomic profiling of the fecal microbiota of mice colonized with the tumorigenic or non-tumorigenic cultured microbiota collections across multiple mice and timepoints revealed mostly commensal populations dominated by *Bacteroides, Bifidobacterium, Clostridium*, and *Escherichia* (Figure 1D and S1F). Mice colonized with the tumorigenic HD2 microbiota saw a shift from *Bifidobacterium, Coprococcus* and *Dorea* to *Clostridium* and *Enterococcus* over time perhaps due to inflammation. Except for these changes, intestinal microbial composition and diversity remained largely stable and donor-specific across time (Figure S1E), suggesting tumorigenesis was mediated through the complex microbiome present at the time of colonization and is not dramatically altered by early tumorigenesis given that these mice infrequently yield colon carcinomas.

### Genotoxic, but not pro-inflammatory or pro-proliferative, bacteria induce tumorigenesis in *Apc*^*Min/+*^;*Il10*^−/−^ mice

Given the emerging evidence suggesting bacterial strains influence tumorigenesis through their ability to drive inflammatory, proliferative, or genotoxic phenotypes, we used high-throughput in vitro screens to identify strains from the tumorigenic HD2 microbiota that functionally alter each of these properties in mammalian cell lines. To identify inflammatory strains, we explored our previously published work detailing cytokine responses of bone marrow derived dendritic cells (BMDCs) co-cultured with filtered conditioned media from 48-hour bacterial cultures^42^. We characterized pro-inflammatory strains based on their ability to induce TNFα, an important pro-inflammatory mediator that has been implicated in the progression of metastatic CRC^43,44^. We found BMDC responses to bacteria in the HD2 community led to a broad range of TNFα secretion that significantly varied across strains (p<1 x 10^-4^, ANOVA; Figure 2A). To assess whether TNFα-inducing bacterial strains lead to tumorigenesis, we colonized GF *Apc*^*Min/+*^;*Il10*^−/−^ mice with a pool of bacterial strains from the tumorigenic HD2 microbiota that induced TNFα levels above the mean TNFα in vitro (“HD2-TNFα+”). We found that high-TNFα-inducing bacteria did not lead to tumors in our CRC-susceptible mice (Figure 2D), suggesting that our in vitro innate immunogenicity screen is not predictive of tumor-promoting microbes in *Apc*^*Min/+*^;*Il10*^−/−^ mice.

**Fig. 2:**
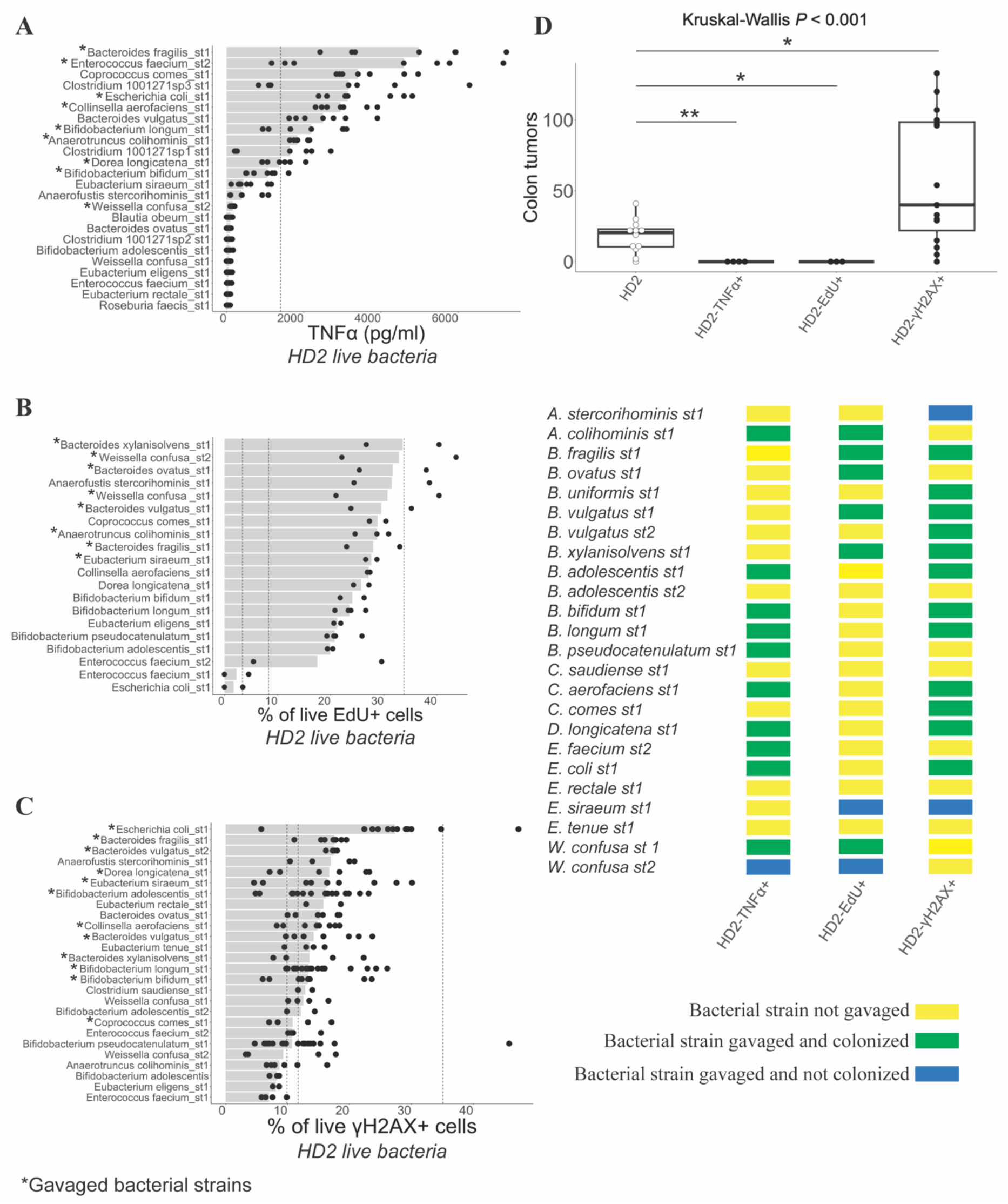
Genotoxic, but not pro-inflammatory or pro-proliferative, bacteria induce tumorigenesis in *Apc*^*Min/+*^;*Il10*^−/−^ mice. **(A)** TNFα levels as measured by ELISA from BDMCs co-cultured with the live bacteria from the HD2 microbiota (presented in rank order). Each bar represents the median concentration of TNFα induced by a particular condition. Dashed vertical line represents the mean TNFα response to a common enteric pathogen, Salmonella enterica (1400 pg/ml), included for reference. Asterisk next to bacteria on the y-axis indicate gavaged strains. **(B)** Proportion (% of live cells) of EdU staining of HT-29 cells co-cultured with the live bacteria from the HD2 microbiota (presented in rank order). Each bar represents the median proportion of EdU incorporation induced by a particular condition. Dashed vertical line represents the proportion of EdU+ HT-29 cells treated with 1000nM staurosporine (ST) (3.3%), 500nM ST (8.3%), or 500ng/ml epidermal growth factor (EGF) (33.8%) (from left to right). **(C)** Proportion (% of live cells) of γH2AX in HT-29 cells co-cultured with the live bacteria from the HD2 microbiota (presented in rank order). Each bar represents the median proportion of γH2AX induced by a particular condition. Dashed vertical lines represent the mean γH2A.X elicited by the live E. coli NC101 (35.1%), E. coli MG1655 (11.7%), and ETBF (9.9%). **(D)** Colon tumor number in GF Apc^Min/+^;Il10^−/−^ mice colonized with a pool of TNFα-(n=4), EdU-(n=3), or γH2A.X-inducing (n=4) bacteria in the HD2 community. The list of bacterial strains represents the 24 unique strains present in the complete HD2 community. B. uniformis_st1 and B. adolescentis_st1 failed to engraft in the whole community but did engraft with the HD2-γH2AX+ subset. The yellow box under each of the HD2 subsets indicates that the strain was not gavaged in the mouse; the green box indicates that the strain was gavaged and had engrafted; blue indicates that the strain was gavaged but had not engrafted in the mouse. The presence or absence of a bacterial strain in a mouse was confirmed by metagenomics. Each point represents data from one mouse. Statistical significance was calculated using Kruskal– Wallis followed by Mann–Whitney tests (**p< 0.01, *p< 0.05). The HD2 (n=12) colon tumor data re-plotted from Figure 1B are shown as white points.

We next sought to identify bacteria that increase epithelial cell proliferation, as cancer-associated bacteria have been shown to alter signaling pathways involved in cell stemness and growth^45 46^. To assess the influence of bacteria on cell proliferation, we co-cultured HT-29 cells with the live bacteria or the stationary phase conditioned media of each bacterial strain from the HD2 community in the presence of EdU (5-ethynyl-2’-deoxyuridine), a nucleoside analog of thymidine. We then measured the incorporated EdU using a fluorophore-labeled azide in a copper-catalyzed click chemistry reaction using flow cytometry. We found a broad range of proliferative capacity across the HD2 bacterial strains (Figure 2B and S2A). EdU incorporation, reflecting DNA synthesis, was highest in cells co-cultured with two specific *Weissella confusa* strains. To determine the tumorigenic potential of pro-proliferative bacterial strains in this donor library, we colonized GF *Apc*^*Min/+*^;*Il10*^−/−^ mice with bacterial strains inducing EdU incorporation above the mean across both assays (“HD2-EdU+”). This subset failed to induce tumorigenesis in our mice at 16 weeks post-colonization (Figure 2D) suggesting that the pro-proliferative bacterial strains in the context of HT-29 cells are not tumorigenic in the *Apc*^*Min/+*^;*Il10*^−/−^ model.

Finally, we explored genotoxicity as the potentially causative microbe-driven mechanism in colorectal carcinogenesis given prior evidence of genotoxic microbes promoting tumorigenesis^6,16,33,34,39,47^. To assess strain genotoxicity, we co-cultured HT-29 cells with the live bacteria or the stationary phase conditioned media from the HD2 bacteria cultures. We then quantified DNA damage using a flow cytometry-based detection of histone variant H2A.X phosphorylated on serine 139 (γH2AX), an important checkpoint for homologous recombination and non-homologous end joining repair pathways and thus, a marker of DNA double-strand breaks (DSBs) (Figure 2 and S2)^48,49^. Colibactin-producing *E. coli* strain NC101, previously established to increase γH2AX in HT-29 cells^17^ and induce tumorigenesis in *Apc*^*Min/+*^ mice^10^, and non-colibactin producing strain, MG1655, served as positive and negative controls, respectively. ETBF served as an additional positive control^37^. We observed significant variation in genotoxicity across strains in the tumorigenic HD2 microbiota with both the live bacteria and the stationary phase conditioned media assays (p=1.28 x 10^-5^ and p=6.70 x 10^-8^ respectively; ANOVA; Figure 2C and S2B), with no correlation between the DNA-damage-inducing strains and assay type (p=0.290; Pearson; live bacteria vs conditioned media). Across the three in vitro screens, only *B. fragilis* induced high levels of TNFα, EdU, and γH2AX. We colonized GF *Apc*^*Min/+*^;*Il10*^−/−^ mice with a pool of HD2 bacterial strains that were above the mean γH2AX in either or both assays (“HD2-γH2AX+”). This cocktail of genotoxic bacteria led to a significantly greater number of tumors than even the complete tumorigenic HD2 community (p=0.023; Figure 2D). These data suggest that genotoxicity is predictive of tumorigenic bacteria in our animal model. In addition, the higher tumor counts in the HD2-γH2AX+ mice relative to the complete HD2 community raises the notion that communities might be composed of a balance of pro-tumorigenic and anti-tumorigenic strains that ultimately determine tumor counts.

### Genotoxic microbes are common in human gut microbiotas

We expanded the γH2AX screening to 304 unique bacterial strains from the tumorigenic (N=6) and non-tumorigenic (N=6) donor microbiotas we screened in vivo (Figure 1B; Figure S3A and S3B). For reference, we also included known genotoxic or tumor-associated species *M. morganii*^39^, *B. fragilis*^37^ and *E. coli* isolated from additional donors^50^, with *E. coli* strains NC101 and MG1655 again serving as a positive and negative control, respectively. We found a tremendous range of genotoxic potential that significantly varied across strains by several orders of magnitude in the live bacteria and the stationary phase conditioned media screens (both p<1 x 10^-4^, ANOVA; Figure 3A and 3B), suggesting the widespread nature of gut microbiota-mediated genotoxicity. We defined “genotoxic” bacteria as those inducing γH2AX above the mean + two times the standard error of the mean (SEM). As expected, we observed an increase in γH2AX phosphorylation in cells infected with the live *E. coli* NC101 strain (Figure 3A)^51,52^. This effect was not observed with the cell-free conditioned media of *E. coli* NC101 culture due to the high instability of colibactin^53,54^ (Figure 3B). Conversely, conditioned media from ETBF induced increased γH2AX, whereas co-incubation with the live ETBF failed to induce γH2AX (Figure 3A and 3B). *E. coli* MG1655 or medium alone did not induce substantial DNA damage across both assays (Figure 3A and 3B). The DNA damage responses to these known genotoxic or tumor-associated species largely fell in the lower to middle part of the γH2AX distributions, suggesting that γH2AX induction by many of commensal isolates is on average higher than that of known pathogens (i.e., *E. coli* NC101, ETBF). The exception to this observation is *E. coli* NC101 in the live bacteria screen, which induced substantial DNA damage with only a few other species from CRC, IBD, and HDs eliciting comparable genotoxicity (Figure 3A; *K. pneumoniae, E. faecium, E. avium, C. koseri*, and other *pks*^+^ *E. coli*). Together, these results indicate that DNA damage responses induced by gut-derived bacterial strains span a broad dynamic range and can be as strong as responses elicited by well-characterized tumorigenic bacteria (i.e., *E. coli* NC101, ETBF).

**Fig. 3:**
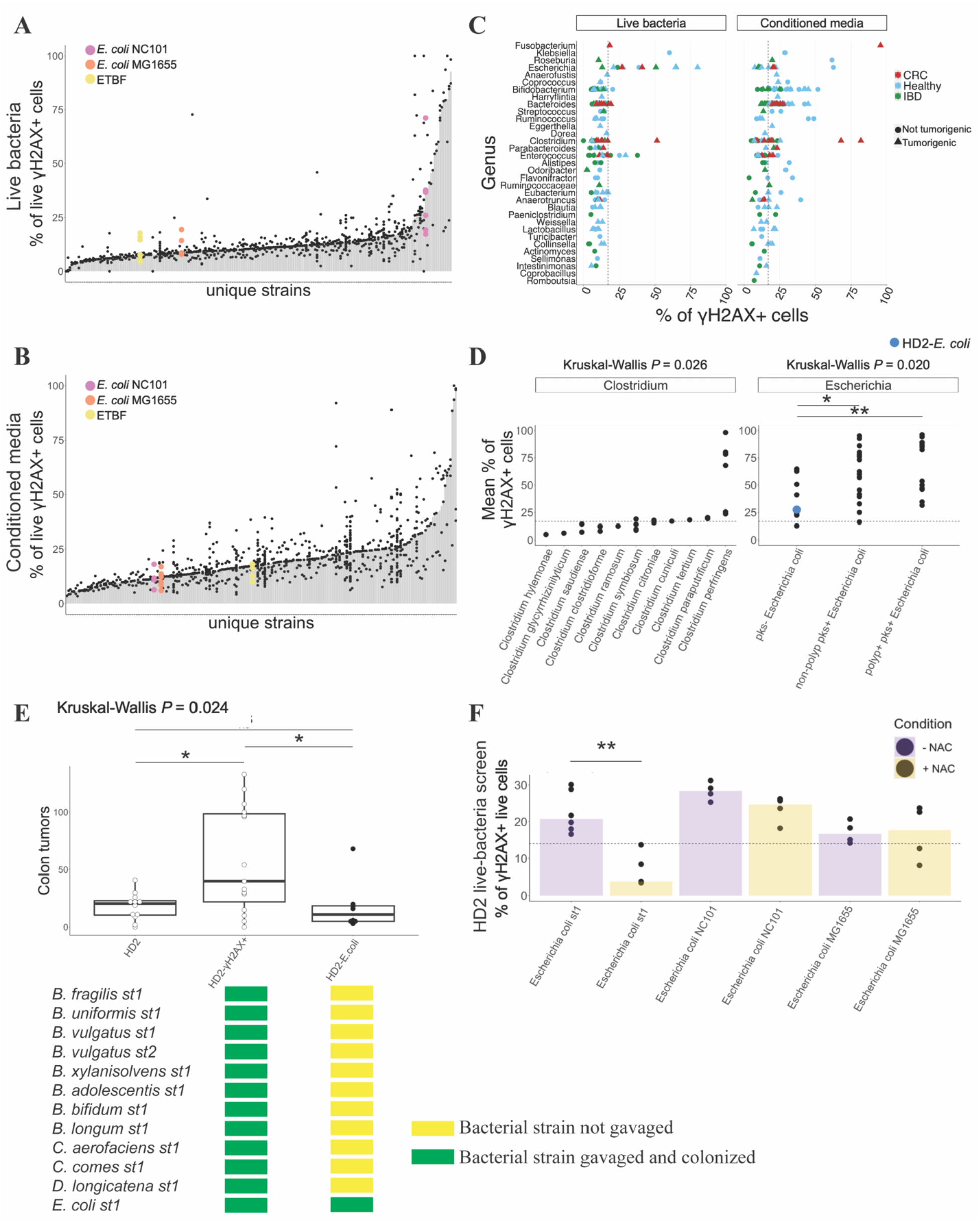
Genotoxic microbes are common in human gut microbiotas. **(A)** Proportion (% of live cells) of γH2AX in HT-29 cells co-cultured with the live bacteria, or **(B)** the stationary phase conditioned media in the 304-strain library (presented in rank order). Each bar represents the median proportion of γH2AX induced by a particular condition. E. coli NC101 (magenta), E. coli MG1655 (orange), and ETBF (yellow) are included for reference. **(C)** Proportion (% of live cells) of γH2AX in HT-29 cells co-cultured with the live bacteria (left), or the stationary phase conditioned media (right) from the 304-strain master library cultures (presented in rank order across both screens). Black dashed vertical line represents the genotoxicity threshold determined by mean + 2 x SEM γH2AX induced by HT-29 cell media alone (16.9%). Each point represents the mean value from a group of 2-6 replicates. The color and shape of each point represents the health status of the originating donor and the in vivo tumorigenic profile, respectively. **(D)** Mean proportion (% of live cells) of γH2AX in HT-29 cells co-cultured with either the live bacteria or the stationary phase conditioned media from the 304-strain library cultures across the two of the four most common genera, with an additional 23 pks^+^ E. coli strains isolated from individuals with or without polyps. Black dashed horizontal line represents the genotoxicity threshold determined by mean + 2 x SEM γH2AX induced by HT-29 cell media alone (16.9%). Each point represents the mean value from a group of 2-6 replicates selected from either screen type depending on the which screen had the maximum value. The E. coli strain from the HD2 library is highlighted in blue. **(E)** Macroscopic colon tumors in gnotobiotic Apc^Min/+^;Il10^−/−^ mice colonized with the γH2AX-inducing E. coli (n=8) from the HD2 microbiota. HD2 (n=12) and HD2-γH2AX+ (n=15) colon tumor data re-plotted from Figure 1B and 2D are shown as white points. Tumors counted at 16 weeks post-colonization. Each point represents data from one mouse. The list of bacterial strains represents the 12 γH2AX-inducing strains in the complete HD2 community. The yellow box under each of the HD2 subsets indicates that the strain was not gavaged in the mouse; the green box indicates that the strain was gavaged and had engrafted. The presence or absence of a bacterial strain in a mouse was confirmed by metagenomics. **(F)** Proportion (% of live cells) of γH2AX in HT-29 cells co-cultured with the live pks^-^ E. coli from the HD2 microbiota in the presence or absence of 20mM NAC. E. coli NC101 and E. coli MG1655 are included for reference. Statistical significance was calculated using Kruskal–Wallis followed by Mann-Whitney tests (**p< 0.01, *p< 0.05, ns: not significant).

To examine if specific bacterial taxonomic groups were associated with the induction of γH2AX, we looked at genera within the 304-strain library to explain the variability associated with genotoxicity. We found statistically significant differences in DNA-damaging activity across genera in the live bacteria and the stationary phase conditioned media screens (both p<1 x 10^-4^; ANOVA; Figure 3C). Specifically, bacteria belonging to the *Clostridium* (gram-positive) and *Escherichia* (gram-negative) genera induced substantial γH2AX in the live bacteria screen. In the stationary phase conditioned media screen, genotoxic bacteria largely belonged to the *Bacteroides* (gram-negative), *Bifidobacterium* (gram-positive), and *Clostridium* genera.

Although previously described genotoxins (e.g., cytolethal distending toxin or CDT, BFT, colibactin) are primarily produced by gram-negative bacteria including *E. coli* and *B. fragilis*, we found multiple gram-positive bacteria (i.e., *B. adolescentis, C. perfringens*) induced substantial DNA damage. DNA-damaging activity was largely independent of donor’s health status as both CRC- and non-CRC cultured communities contained genotoxic microbes.

Having identified broad taxa with highly genotoxic bacteria, we next focused on the four genera with the highest number of strains per genus in our library and highly varied γH2AX induction either in the live-bacteria or the stationary phase conditioned media screen (*Bacteroides* = 51 strains, *Bifidobacterium* = 25 strains, *Clostridium* = 20 strains, and *Escherichia* = 45 strains). We observed that the *Clostridium* members exhibited strong DNA-damaging activities, with the two unique *C. perfringens* strains derived from CRC patients among those which most highly induced γH2AX (p=0.026, Kruskal-Wallis; Figure 3D). There were no significant differences in the γH2AX induction across the different species of *Bacteroides* or *Bifidobacterium* (Figure S3C). *E. coli* exhibited a broad range of genotoxicity (9.2% - 96.1% γH2AX+ cells) that significantly varied by strain (p=0.020, Kruskal-Wallis; Figure 3D). Given the established role of *pks* harboring *E. coli* in tumorigenesis and the paucity of *pks*^*+*^ *E. coli* isolates in our original 304-strain library, we screened an additional 23 *pks*^+^ strains isolated from individuals with or without polyps. We observed a significant increase in the genotoxicity of *pks*^+^ *E. coli* isolated from polyps or non-polyps relative to *pks*^-^ *E. coli* (p=0.008, p=0.04; respectively; Figure 3D). Within the *pks*^+^ *E. coli* strains, presence of donor polyps did not predict a difference in γH2AX induction (p=0.399). All 12 original subjects whose microbes were assessed in the screen harbored one or more genotoxic bacteria, highlighting the importance of assessing genotoxicity of strains regardless of donor health status.

### Genotoxic *pks*^-^ *E. coli* induces tumorigenesis and elicits DNA damage in vitro through ROS production

As previously mentioned (Figure 2D), the HD2-derived genotoxic consortium containing 14 unique bacterial strains (12 of which engrafted in vivo) led to a significantly greater number of colon tumors than the complete HD2 microbiota, suggesting we had tipped the balance more in favor of tumorigenesis by colonizing the mice with tumorigenic bacteria and removing protective bacteria. To explore individual versus group effects of these strains in vivo, we colonized GF *Apc*^*Min/+*^;*Il10*^−/−^ mice with the genotoxic *pks*^-^ *E. coli* strain from the HD2 microbiota that consistently induced substantial γH2AX in both assay screens (Figure 3E). The HD2-*E. coli* monocolonization led to a similar number of tumors as the complete HD2 community (p=0.374) but induced significantly fewer tumors than the complete HD2-γH2AX+ subset (p=0.036). These results suggest that *E. coli* is one of the drivers of tumorigenesis by the HD2 microbiota, and co-colonization with additional genotoxic bacteria is required for maximum tumor burden in our animal model – further reinforcing the concept of cooperation among multiple pathogenic bacteria in compounding tumor burden.

Notably, the potent γH2AX-inducing *E. coli* strain in the HD2 microbiota does not harbor genes encoding colibactin or CDT, suggesting additional genotoxins remain to be characterized. We confirmed the absence of colibactin production by the HD2-*E. coli* using PCR primer pairs targeting either end of the *pks* pathogenicity island^17^, as well as using screens quantifying NMDA^55,56^, the most abundant colibactin intermediary product (Figure S3D). We similarly confirmed the absence of CDT production by the HD2-*E. coli* using PCR primer pairs targeting the bioactive CDT subunit, *cdtB*^57^ (Figure S3E). Moreover, we determined that γH2AX induced by the genotoxic HD2 *pks*^-^ *E. coli* is significantly diminished in the presence of an antioxidant, N-acetylcysteine (NAC), suggesting a ROS-mediated DNA damage mechanism (p=0.004, Figure 3F). Gene set enrichment analyses of differentially expressed genes from RNA-seq of HT29 cells co-cultured with the stationary phase conditioned media of the HD2-*E. coli* culture revealed an increased expression of genes involved in redox metabolism (Figure S3F), further supporting the mechanism of DNA DSBs by the HD2-*E. coli* through the generation of ROS.

### Genotoxic human gut microbes have an additive effect on tumor burden and accelerate tumorigenesis in susceptible mice

To determine if the increased tumorigenicity of the genotoxic bacterial subset relative to the complete community generalizes to other human gut microbiota communities screened for the γH2AX induction (Figure 3), we examined a second tumorigenic microbiota from a CRC donor. We observed a similar magnitude increase, although not statistically significant, in colon tumor counts of mice colonized with a pool of genotoxic bacterial strains from the CRC1 community (mean tumor counts CRC1 community = 19.3 vs mean tumor counts CRC1-γH2AX+ = 57.3; Figure 4A). A pool of non-genotoxic bacterial strains from the CRC1 community (CRC1-γH2AX-) led to a significantly lower number of tumors than the CRC1-γH2AX + consortium (p=0.005; Figure 4A, S4A and S4C). To explore if non-tumorigenic microbiotas (Figure 1B) might harbor tumorigenic microbes whose pathogenic impact is more completely hampered by protective microbes, we colonized GF *Apc*^*Min/+*^;*Il10*^−/−^ mice with the pool of γH2AX+ bacteria from the non-tumorigenic HD7 donor library. We found a significant increase in tumor burden in mice colonized with the HD7-γH2AX+ bacterial pool compared to the complete HD7 community (p=0.024; Figure 4A, S4B and S4D). Together these results further support our earlier observation (Figure 2D) that the combined impact of tumorigenic and protective microbes determines the in vivo tumor count.

**Fig. 4:**
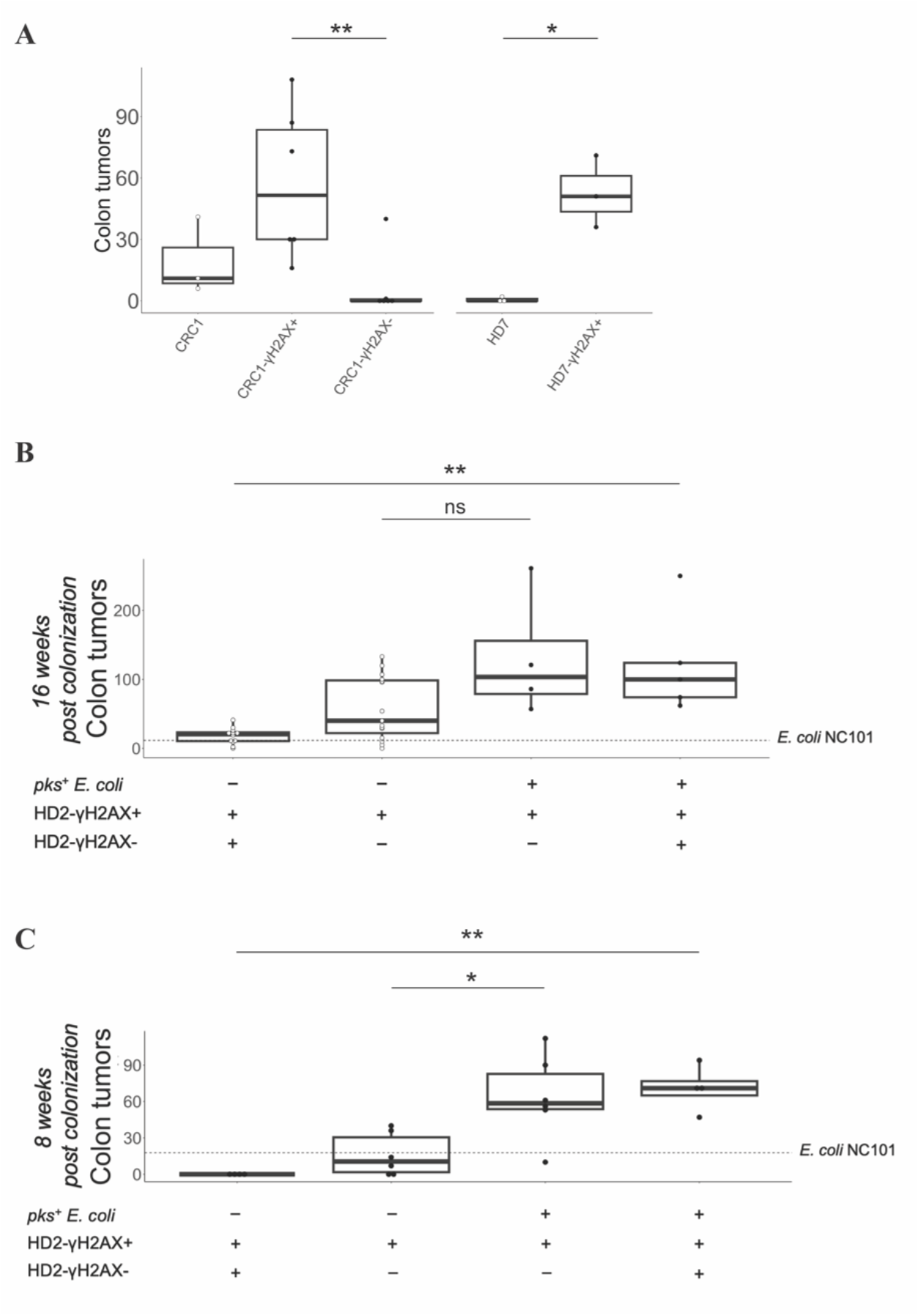
Human gut microbiotas have a cooperative effect on tumor burden and accelerate tumorigenesis in susceptible mice. **(A)** Colon tumor numbers in GF Apc^Min/+^;Il10^−/−^ mice colonized with a pool of γH2A.X-inducing (n=6) or non-γH2A.X-inducing (n=6) bacteria from the tumorigenic CRC1 microbiota or with a pool of γH2A.X-inducing (n=3) bacteria from the non-tumorigenic HD7 microbiota. Dunn’s unadjusted P values shown (*p< 0.05, **p< 0.01). **(B)** Macroscopic colon tumors detected in HD2 mono-colonized (n=12), HD2-γH2A.X+ mono-colonized (n=15), E. coli NC101/HD2-γH2A.X+ co-colonized (n=4), or E. coli NC101/HD2 co-colonized (n=5) Apc^Min/+^;Il10^−/−^ mice at 16 weeks post-colonization. Dashed horizontal line represents the mean colon tumor counts detected in E. coli NC101 mono-colonized (n=6) mice at 16 weeks post-colonization. **(C)** Macroscopic colon tumors detected in HD2 mono-colonized (n=4), HD2-γH2A.X+ mono-colonized (n=6), E. coli NC101/HD2-γH2A.X+ co-colonized (n=6), or E. coli NC101/HD2 co-colonized (n=4) at 8 weeks post-colonization. Dashed horizontal line represents the mean colon tumor counts detected in E. coli NC101 mono-colonized (n=5) mice at 8 weeks post-colonization. Each point represents data from one mouse. Statistical significance was calculated using Kruskal–Wallis combined with Dunn’s multiple pairwise comparisons test with BH adjustment (*p< 0.05, **p< 0.01, ns: not significant). Colon tumor data re-plotted from prior figures are shown as white points.

To further explore the bacteria-mediated cooperative effects on tumorigenesis, we colonized mice with the tumorigenic HD2 microbiota in combination with the known tumorigenic *pks*^+^ *E. coli* murine strain NC101. The combination of these bacteria led to a robust tumorigenesis that far exceeded tumors induced by HD2 microbiota alone or the *E. coli* NC101 alone (p=0.005 and p=0.007, respectively; Figure 4B) further demonstrating that increasing the balance of the microbiota towards pro-tumorigenic bacteria increases tumor count.

We next sought to determine whether colonization with multiple pathogenic strains could also accelerate tumor onset, we colonized GF *Apc*^*Min/+*^;*Il10*^−/−^ mice with *E. coli* NC101 or the HD2 microbiota separately or together and analyzed intestinal tissues for macroscopic tumors at 8 weeks post-colonization, instead of 16 weeks. We found that at 8 weeks post-colonization, the complete HD2 microbiota alone did not lead to colon tumor formations in the *Apc*^*Min/+*^;*Il10*^−/−^ mice. However, in combination with *E. coli* NC101, the HD2 microbiota induced substantially greater number of colon tumors relative to the HD2 microbiota alone (p=0.009; Figure 4C) demonstrating a faster tumor onset and suggesting that increasing the load of tumorigenic bacteria can increase tumor number and decrease time to tumor onset. The combination of *E. coli* NC101 and the HD2 microbiota led to a similar magnitude increase, although not statistically significant, in colon tumor counts as *E. coli* NC101 alone (mean tumor counts *E. coli* NC101 + HD2 = 70.8 vs mean tumor counts *E. coli* NC101 alone = 17.8; Figure 4C).

We observed comparable findings in tumor multiplicity and tumor onset when combining the genotoxic subset of HD2 and *E. coli* NC101 (Figure 4C). At 8 weeks post-colonization, the combination of these bacteria led to a substantial increase in colon tumor counts as *E. coli* NC101 (p=0.044; Figure 4C).

## DISCUSSION

In this study, we screened bacterial strains for their ability to promote inflammation, proliferation, and genotoxicity in human colorectal adenocarcinoma cells to better elucidate causal links between gut microbes and disease state. We found the number of tumors in the colon of our mice varied significantly by donor microbiota, regardless of health status of the originating donor. We found that in vitro genotoxicity of bacterial strains predicted in vivo tumorigenesis, while pro-proliferative and innate immunogenicity of strains did not predict tumor counts. We identified a genotoxic and tumorigenic *pks*^-^ *E. coli* whose genotoxicity was neutralized by NAC, suggesting a ROS-mediated mechanism of DNA damage.

Although the gut microbiota has long been implicated in colorectal carcinogenesis, this complex relationship is not well-characterized. Studies describing the effects of the human microbiota and tumorigenesis are often based on a small number of human microbiotas or a few model bacterial strains^40,58^. Our work expands upon the few known model tumorigenic strains as well as a recent genotoxicity screen of phylogenetically diverse human gut commensal bacterial strains from IBD patients^39^. Importantly, we found pools of genotoxic bacterial strains from tumorigenic microbiotas, as well as a non-tumorigenic microbiota, drove greater number of tumors than the complex microbiotas in vivo, while genotoxic-low bacterial strains failed to induce tumorigenesis. These data suggest that the final tumor count is a reflection of a combination of pro-tumorigenic strains and protective strains that inhibit tumorigenesis directly or by reducing the relative abundance of tumorigenic organisms. These results build on previous observations of positive and negative cooperative interactions among microbes in the context of infection^59,60^, cancer^37,61,62^, and anticancer therapies^63,64^. We further confirmed the presence of genotoxic bacteria across 100% of the CRC, IBD, and HDs (N=12) screened in our 304-strain library, supporting the notion that we all harbor microbes that influence the risk of developing CRC, but the combined risk across all microbes in a microbiome varies across individuals.

Recently, the alarming increase in early-onset CRC burden led to new recommendations by the US Preventive Services Task Force initiating CRC screening at age 45 years, reduced from age 50 years in previous versions^65^. The pace of this increase in early-onset CRC implicates environmental factors such as the gut microbiota as an important contributor^3,66^. Our results, as well as earlier results^37^, suggest one mechanism through which the microbiota could influence the onset of CRC, as combining multiple tumorigenic or genotoxic gut microbiotas had an additive effect on tumor burden in our CRC-susceptible mice and reduced the time to tumorigenesis in half. If changes in diet, lifestyle, or antibiotic usage were to alter the balance of tumorigenic to protective bacterial strains harbored in humans, it could drive a population-level reduction in the time to onset for CRC^67–69^.

The work presented here reveals the existence of many pathobionts and previously unexplored gut-derived genotoxic bacteria across IBD and CRC patients, as well as in healthy individuals. Characterizing this understudied majority of human microbes and expanding our functional and mechanistic understanding of the diversity of the gut microbiota could enable new insights into diseases with unknown etiology, provide disease-predictive microbiome signatures, and advance microbial therapeutics or dietary interventions to restore a tumor-protective balance of microbes in the gut.

## MATERIALS AND METHODS

### Animals

The Icahn School of Medicine at Mount Sinai Institutional Animal Care and Use Committee (IACUC) approved all animal experiments (Protocol # 2013-1385 and # 2017-0297). GF 129/SvEv *Apc*^*Min/+*^;*Il10*^−/−^ were bred in flexible vinyl isolators at the Mount Sinai Precision Immunology Institute Gnotobiotic Facility. After weaning, GF mice were aseptically transferred outside of the breeding isolators and colonized with human microbiotas. Following colonization, mice were housed in individually-ventilated, positive pressure cages (Allentown) and handled using strict aseptic technique. All mice were provided sterile water and chow (5K67, LabDiet) and maintained under a strict 12-hour light cycle. The mice were age- and sex-matched when possible.

### Human samples and bacterial cultures

Arrayed culture collections of human gut microbiome-derived bacteria used in this study were previously isolated from donor stools as previously described^24,50^. Fecal samples from donors with early-stage primary tumors in the colon or rectum were obtained from a contract research organization (BioIVT). Samples were collected before chemotherapy or surgery. No donor had taken antibiotics in the three months before sample collection. Of the four donors used in this study, three were diagnosed with cancer of the colon and one with rectal cancer. All donors were assessed as having tumor grade 2 disease, three with T3N0M0 disease and one with T3N0M9 disease. The average age of these donors was 67 +/-3 years. Three were male and one female and all self-identified as Caucasian. These donors had no diagnosis of autoimmune or inflammatory disease, renal or liver disease, dementia, significant mental illness, or hematological/oncological disease (except CRC).

Briefly, donor stool was clarified and plated onto a variety of solid selective and non-selective media under anaerobic, micro-aerophilic and aerobic conditions. After 48-72 hours at 37°C, 384 single colonies from each donor were individually picked and regrown in liquid LYBHIv4 media (37 g/l Brain Heart Infusion [BD], 5g/l yeast extract [BD], 1 g/l each of D-xylose, D-fructose, D-galactose, cellobiose, maltose, sucrose, 0.5 g/l N-acetylglucosamine, 0.5 g/l L-arabinose, 0.5 g/l L-cysteine, 1g/l malic acid, 2 g/l sodium sulfate, 0.05% Tween 80, 20 mg/mL menadione, 5 mg/l hemin (as histidine-hemitin), 0.1 M MOPS, pH 7.2) for 48 hours under anaerobic conditions. Regrown isolates were identified and re-arrayed into 96 well plates (1 bacteria per well, 1 plate per individual). MALDI-TOF mass spectrometry (Bruker Biotyper) and 16S rDNA amplicon sequencing were used to identify individual isolates at the species level, or at the strain level using whole genome sequencing^24,70^. For administration to mice, all regrown isolates were pooled into a single cocktail and stored in LYHBHIv4 media with 15% glycerol.

### Gnotobiotic mouse experiments

4-6 week GF *Apc*^*Min/+*^;*Il10*^−/−^ mice were colonized with 200ul of pooled cocktail of cultured strains by a single oral gavage. Following colonization, mice were housed in autoclaved, filter-top cages outside of the breeding isolators as described above. Mice were colonized for 16 weeks before analysis of colon tumor counts and histological evaluation of intestinal inflammation.

Every two weeks, mice were weighed using a portable precision balance that was disinfected between cages and fecal pellets were collected and stored at −20°C until analysis. Loss of body mass and elevation in fecal LCN2 were used as biomarkers of intestinal inflammation severity. Additionally, fecal pellets were used to determine gut microbiota composition. Mice were sacrificed at specified time-points unless otherwise noted for humane reasons related to poor health and/or excessive weight loss (defined as ≥20% total body weight). We also assessed for sexual dimorphism in colorectal tumorigenesis in the *Apc*^*Min/+*^*;Il10*^−/−^ mice and did not find a significant difference in the number of tumors across male and female mice, although female mice generally exhibited higher tumor burden (Figure S5A).

### Colon Collection and Histology

Mice were euthanized at 16 weeks post-colonization. The small intestine and colon were harvested, cut open longitudinally, and macroscopic tumors counted. The intestine tissues were then Swiss rolled and fixed in 10% neutral buffered formalin solution for 24 hours before transfer to 70% ethanol. Swiss rolls were processed, paraffin-embedded, sectioned (4μm), and Hematoxylin and Eosin (H&E) stained by Histowiz Inc (home.histowiz.com). Slides were scanned to create whole slide image and histological scoring of inflammation was performed blindly by a board-certified gastrointestinal pathologist. Eight histological components were assessed: inflammatory infiltrate, goblet cell loss, hyperplasia, crypt density, muscle thickness, submucosal infiltration, ulceration, and crypt abscesses (all scored from 0-4).

### Tumor quantification

At 8 or 16 weeks post-colonization, mice were sacrificed, and macroscopic tumors were counted in the small intestine and colon using a vernier caliper. Three methodologies for quantifying colon tumor burden were initially compared: 1) macroscopic counts, 2) percent of whole colon, and 3) percent of whole Swiss roll. We used four mice colonized with our most tumorigenic microbial subset for these comparisons (Figure 2D). GF mice were used as a negative control, as *Apc*^*Min/+*^;*Il10*^−/−^ mice are notable for their microbiota-dependent susceptibility to spontaneous tumors in the colon^10^.

For macroscopic counts, tumors were counted at the time of intestinal tissue dissection using a 10X magnification lens and a vernier caliper. Adenomatous regions were subsequently verified using representative H&E images. The second method, percent of whole colon, was calculated by dividing the area of the region containing tumors by the total area of colon (Figure S5B). To calculate the percent of whole Swiss roll, scanned H&E sections were used to measure the area of tumorigenic regions and the total area of colon before dividing these measures by one another (Figure S5C). QuPath software was used to segment and measure the regions of interest.

We compared the similarity between trends (Figure S5D-E) in tumor counts across mice colonized with the HD2-γH2AX+ subset (Method 1 vs 2, r = 0.96, Method 1 vs 3, r = 0.94, Method 2 vs 3, r = 0.99, Pearson correlation coefficient). Given that we saw a strong positive correlation between the methodologies, we chose to proceed with the macroscopically tumor counting method that was the most efficient.

### Lipocalin2

Fresh fecal pellets were collected in sterile, pre-weighed, barcoded matrix tubes and stored in −20°C until analysis. After collection, the stored pellets were weighed to determine the weight of the stool. The fecal pellet was then mixed with sterile PBS in a volume (in uL) that corresponded to 10 times the pellet weight and agitated using Bead-Beater (without any beads present) and Multi-Tube Vortexer for a total 20 minutes. The resulting mixture was then centrifuged at 4000rpm for 20 minutes and supernatant extracted for LCN2 quantification using LCN2 Mouse ELISA kit (R&D systems) according to manufacturer’s instructions. For controls, we utilized stool derived from *Apc*^*Min/+*^;*Il10*^−/−^ mice colonized with well-established tumor-inducing bacterial strains that have been shown to induce severe intestinal inflammation across multiple animal models of CRC (i.e., *E. coli* NC101, ETBF).

### Fecal sample processing, DNA extraction, and shotgun metagenomic sequencing

Fecal microbiome samples were processed as previously described to assay microbial composition and determine if any microbial abundances correlate with tumor formation^70,71^. Briefly, fecal pellets from mice colonized with tumorigenic and non-tumorigenic donor microbiotas were collected in sterile, pre-weighed and barcoded microtubes. These samples were kept on dry ice and then stored at −20°C until further processing. DNA was extracted by bead beating in phenol:chloroform. Illumina sequencing libraries were constructed using the sonicated DNA, ligation products purified, and enrichment PCR performed with Nextera XT DNA Library Preparation Kit (Illumina). Samples were pooled in equal proportions and size-selected before sequencing with an Illumina HiSeq. Reads from metagenomic samples were trimmed and mapped to the unique regions of bacterial genomes known to potentially form part of the microbiota in our experiments. Analysis of sequencing data was performed using a custom software pipeline in R that calculates and visualizes our engraftment metrics.

### Bacterial screening library assembly

To generate our screening library, we regrew preexisting culture libraries from donors of interest in liquid LYBHIv4 media for 48 hours under anaerobic conditions. Unique bacterial strains were selected from each donor library and re-arrayed into 96-well plates. 304 unique isolates from 7 healthy, 5 IBD, and 5 CRC donors were re-arrayed into a master library. Additional strains of *M. morganii, B. fragilis*, and *E. coli* were included as controls. Further inclusion of 23 *pks*^+^ *E. coli* strains from individuals with or without polyps led to a total of 327 isolates comprising our master library. Growth and identity of the regrown master library was confirmed by OD600nm and MALDI-TOF.

### Cell lines

HT-29 cells were obtained from ATCC. The cell line was maintained in McCoy’s 5A with 2200 mg/L sodium bicarbonate, 10% (v/v) fetal bovine serum, 100 U/ml penicillin, 100 mg/ml streptomycin, and 1.5 mM L-glutamine at 37°C with 5% CO2. The cell line was confirmed to be mycoplasma free before use.

### Commensal screening

HT-29 cells were seeded and grown in 96-well flat-bottom plates at 40,000 cells/well in McCoy’s 5A media without added antibiotics (i.e., penicillin, streptomycin). We then began testing responses of these cells to either the filtered stationary phase conditioned media of each bacterial stain in our master library or the live bacterial cultures washed of metabolites. Bacterial libraries were centrifuged at 4000rpm for 10 minutes. Supernatants were removed and passed through a 0.2μm filter (Pall 8019). Bacterial cultures were washed three times with PBS and resuspended in an equal volume of McCoy’s 5A media without added antibiotics. Live bacteria or the stationary phase conditioned media generated by each isolate was then co-cultured with HT-29 cells at a multiplicity of infection (MOI) of 100:1 (bacterial cells:HT-29 cells) or 1:5 (filtered supernatant:cell culture media). The plates were then incubated at 37°C for a 4-hour live cell infection or a 24-hour supernatant co-culture. HT-29 cells were then washed four times with PBS and resuspended in McCoy’s 5A medium with added antibiotics (100 U/ml penicillin, 100 mg/ml streptomycin, and 200μg/mL of gentamicin (Thermo)) to prevent bacterial growth. The cells were subsequently processed and stained for DNA damage (γH2AX) or proliferation (EdU) markers (Figure 2.3). HT-29 cells initially co-cultured with live bacteria were allowed 20 hours to recover at 37°C after antibiotic treatment. For the initial screen involving the 304-strain library, we performed three independent experiments and included multiple technical replicate conditions per experiment. The bacteria co-cultured with HT29 cells were re-grown anaerobically to confirm sustained viability over the duration of the experiment.

### High-throughput γH2AX detection

DSBs were quantified using the γH2AX Phosphorylation Assay Kit for Flow Cytometry (Millipore, 17-344). HT-29 cells were harvested, pelleted, and stained with the Zombie Aqua Fixable Viability dye (Biolegend) for 30 minutes at 4°C to exclude dead cells from analyses. For analysis of the intracellular γH2AX, HT-29 cells were fixed using a 2% formaldehyde solution for 30 minutes at 4°C. The cells were then washed twice with PBS and permeabilized overnight with 0.3% (v/v) Triton X-100 containing FITC-conjugated γ-Histone H2A.X (serine 139) antibody at [0.0018mg/ml]. Hydrogen peroxide (H_2_O_2_) and N-acetyl-L-cysteine (NAC) were used as positive and negative controls, respectively. Additional controls included colibactin-producing *E. coli* strain NC101 and a commensal non-colibactin-producing *E. coli* strain MG1655. Samples were acquired on a five laser Aurora Spectral Cytometer (Cytek) and data analysis performed in FlowJo X (BD Bioscience; Figure 2.4). The stationary phase conditioned media of the *E. coli* NC101 culture did not induce increased γH2AX; a finding that is in line with previous research demonstrating the requirement for direct cell contact for *E. coli* NC101 to drive DNA damage^17^.

### High-throughput EdU detection

DNA replication (EdU) in proliferative HT-29 cells was measured using the Click-it EdU Assay Kit for Flow Cytometry (Invitrogen, C10636). In brief, HT-29 cells were labeled with 0.05μM of EdU during the 20-hour recovery period with the live bacteria co-cultures, or for the full 24-hour co-culture with the stationary phase conditioned media. Dead cells were excluded from analyses as previously described. The cells were fixed for 30 minutes and permeabilized overnight according to the manufacture’s protocol. EdU was then detected by resuspending the cells in a reaction cocktail of copper protectant and fluorescent dye picolyl azide before acquiring the samples on Aurora Spectral Cytometer. Epidermal growth factor (EGF) and a range of staurosporine (ST) concentrations were used as positive and negative controls, respectively.

### High-throughput TNFα detection

TNFα data was obtained from previously published work detailing cytokine responses of bone marrow derived dendritic cell (BMDC) to 277 bacterial strains isolated from the fecal microbiota of 17 humans^42^. In brief, non-adherent cells were isolated from femurs of C57Bl/6J mice and differentiated in vitro using GM-CSF for 8 days. Differentiated BMDCs were then harvested, washed, plated in 96-well plates at a density of 50,000 cells/well and rested overnight before use. The following day, BMDCs were co-cultured with either the filtered conditioned media of each bacterial stain or the live bacterial culture washed of metabolites. Supernatants were harvested after 24 hours, and the specific secreted inflammatory cytokines measured using high-throughput ELISA in a 384-well format.

### PCR detection of *pks*^+^ or cdtB+ E. coli

All *E. coli* strains were screened for the presence of the *pks* island by PCR using the following primers^17^ that target either end of the *pks* island:

Ec_F: 5’-CAT GCC GCG TGT ATG AAG AA-3’ and Ec_R: 5’-CGG GTA ACG TCA ATG AGC AAA-3’, product size=96pb; pksL1: 5’-AAT CAA CCC AGC TGC AAA TC-3’ and pksL2: 5’-CAC CCC CAT CAT TAA AAA CG-3’, product size = 1824bp; pksR1: 5’-AGC CGT ATC CTG CTC AAA AC-3’ and pksR2: TCG GTA TGT CCG GTT AAA GC-3’, product size = 1413bp.

The *E. coli* strain from the HD2 library was further screened for the presence of the bioactive CDT subunit, cdtB, by PCR using the following primers^57^: F: 5’-TGG GCT AGC AGG AGG AAT TCA TGA AAA AAA TTA TAT GTT TAT TTT TAT CTT TTA ACC TT-3’ and R: 5’-TCA TGG TCT TTG TAG TCC ATC TAA AAT TTT CTA AAA TTT ACT GGA AAA TGA TCT GAA AC-3’, product size=251pb. A synthetic gene construct spanning the primer sites was cloned in a plasmid (Integrated DNA Technologies, IA, USA) and used as a positive control.

Briefly, bacteria were grown anaerobically for 48 hours as described above. DNA was extracted by bead-beating followed by QiaQuick columns (QIAGEN) and quantified by Broad Range Quant-IT dsDNA Assay Kit (Invitrogen) alongside a BioTek Synergy HTX Multi-Mode Reader. PCR was performed on the C1000 Touch PCR thermal cycler (Bio-Rad) using SYBR Safe DNA Gel Stain (Thermo Fisher Scientific). PCR products were visualized on an agarose gel using the Gel Doc EZ Imager (Bio-Rad).

### NMDA Quantification Assay

The concentration of NMDA (N-Myristyl-D-Asperigen) was measured using LC-MS analysis. This analysis was performed on a system comprising a Vanquish UPLC (Thermofisher Scientific Inc) interfaced with an Exploris-240, high-resolution mass spectrometer (Thermofisher Inc). Chromatographic separation was achieved using a Poroshell HPH C18 column (2.7 μm, 2.1 x 50 mm, Agilent) with a mobile phase consisting of 10 mM NH4OAC in water (A) and acetonitrile (B). The mass spectrometer was set to operate in negative ionization mode and selected ion monitoring (SIM) mode, with a resolution of 120,000, tracking m/z 341 and m/z 346 within a ± 5 Dalton scan range. For LC-MS quantification, the supernatant from cell culture was mixed in equal volume of a deuterium-labeled d5-NMDA (IS, 20 ng/mL) that was synthesized at Johnson and Johnson. After vortexing and centrifugation, the mixture was injected into the LC-MS system for analysis. NMDA concentration in the sample was calculated based on the peak area ratio of NMDA to d5-NMDA (341.2446/346.2760) in the sample against the standard curve. The standard curve was established in the blank suspension of cell culture with the linear range 0.2-100 ng/mL.

### Bulk RNA-sequencing

Co-culture samples of bacterial strains with 40000 HT29 cells in 96-well plates were frozen in RNAlater (Thermo AM7020), followed by lysis and RNA purification using RNeasy 96 kit (Qiagen 74181). Total RNA samples were reverse transcribed with Maxima H Minus RT (Thermo EP0751) and oligo-dT primers. Sequencing libraries were prepared using NEBNext Ultra II FS Library Prep Kit (NEB E7805S) and sequenced on an Illumina NovaSeq (PE150nt). FASTQ files were trimmed for adapters using FastQC (0.11.9) and cutadapt (4.2)^72^. STARsolo (2.7.10a) was used to map cDNA sequences to reference genome, tabulate reads per gene for each sample barcode, and deduplicate reads into unique transcripts by UMI^73^. Relative gene expression between each sample and untreated control was calculated using the DESeq2 R package (1.44.0)^74^. Gene set enrichment analysis (GSEA) was done using the fgsea (1.30.0) R package^75^ with log2 fold change vs. control as input. Reference dataset for GSEA was downloaded from Moleular Signatures Database^76^.

### Statistical Analysis

Significance thresholds of ***p< 0.001, **p< 0.01, *p< 0.05, or NS: not significant (p> 0.05) are shown. Statistical analyses for all figures were performed with RStudio v4.3.3 using the Wilcoxon or Mann Whitney nonparametric unpaired two-tailed t-test. For correlational studies, we used the Spearman nonparametric correlation analysis with a confidence interval of 95% and a *P*<0.05 threshold applied for statistical significance with two-sided comparisons. Kruskal– Wallis followed by the Dunn’s multiple comparison test with Benjamini-Hochberg (BH) adjustment was carried out in cases of 3+ groups and small sample size (n=3).

## Acknowledgements

This work was supported in part by the staff and resources of the Icahn School of Medicine Gnotobiotic Facility, the Microbiome Translation Center at Mount Sinai, and the Mount Sinai Flow Cytometry Core. This work was supported in part through the computational and data resources and staff expertise provided by Scientific Computing and Data at the Icahn School of Medicine at Mount Sinai and supported by the Clinical and Translational Science Award (CTSA) grant UL1TR004419 from the National Center for Advancing Translational Sciences. We thank C. Fermin, E. Vazquez, and G.N. Escano for gnotobiotic husbandry. This work was supported by the National Institutes of Health grants (nos. NIDDK DK112978 and NIDDK DK124133), and Janssen Research & Development.

## Data and materials availability

All data used to construct figures are available in the supplementary materials. Materials used in this study are available upon request to the corresponding author. Material transfer agreements may be required.

