## Supplement for "Cooperation Between Genotoxic Bacteria Accelerates Tumorigenesis in a Mouse Model of Colon Carcinogenesis"

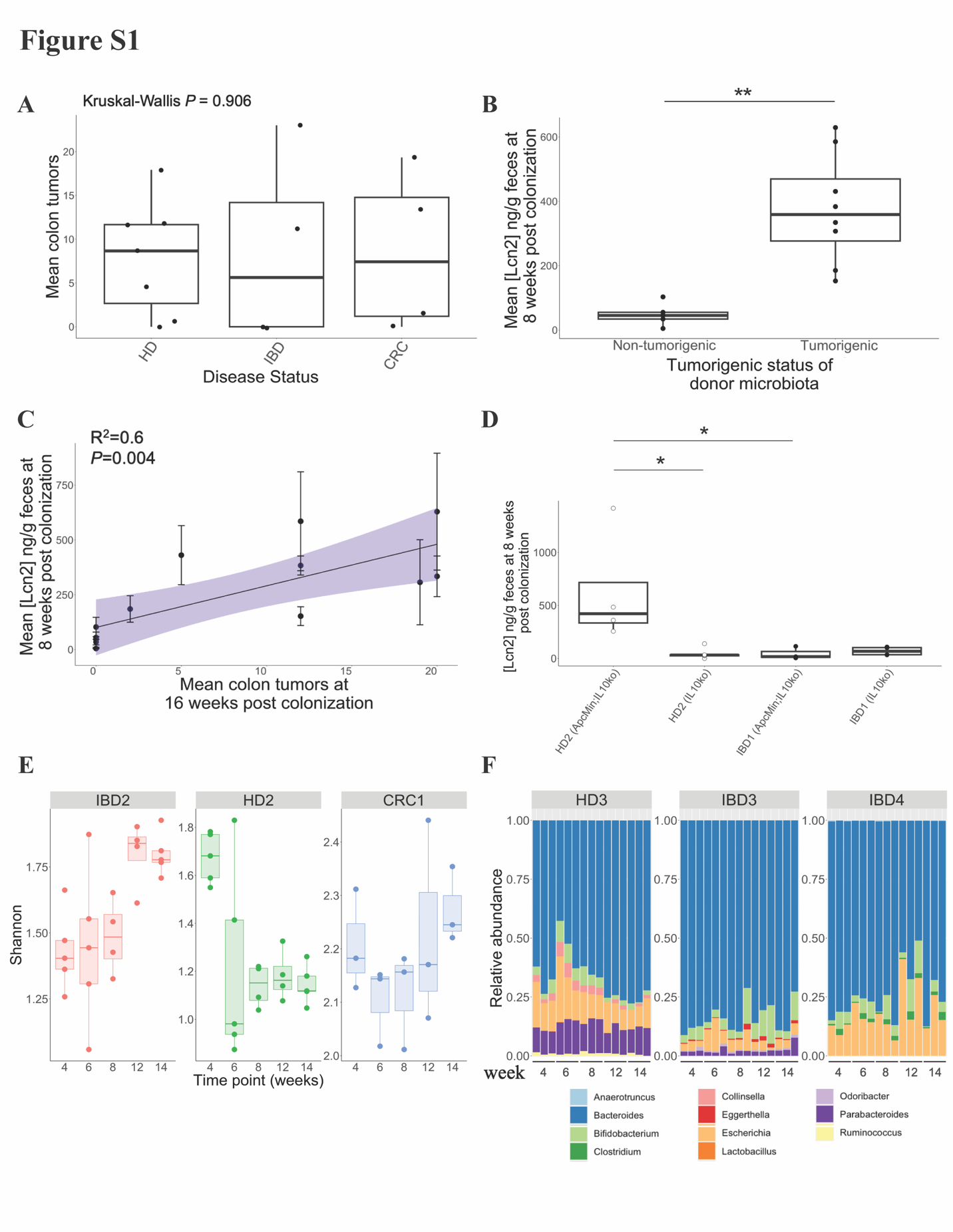

**Fig S1: (A)** Mean colon tumor counts per donor of all microbiomes screened in Figure 1B plotted by disease status (HD=Healthy donor, IBD=Inflammatory bowel disease, CRC=Colorectal cancer). Statistical significance was calculated using Kruskal–Wallis.

**(B)** Mean fecal LCN2 concentrations measured at 8-weeks post*-*colonization in mice gavaged with either non-tumorigenic or tumorigenic microbiota from Figure 1B. Tumorigenic microbiota defined as having a mean tumor count greater than 1.0. Statistical significance was calculated using Kruskal–Wallis followed by Mann–Whitney tests (**p< 0.01). **(C)** Correlation between the mean LCN2 concentrations at 8 weeks post*-*colonization and mean colon tumors at 16 weeks post*-*colonization in mice colonized with the same microbiota. The purple region indicates the 95% confidence interval. P values calculated by F-test. **(D)** Macroscopic colon tumors in *Apc^Min/+^*;*Il10^-/-^* and *Il10^-/-^* mice colonized with tumorigenic HD2 microbiota or non-tumorigenic IBD1 microbiota. Mice were colonized for 8 weeks. The *Apc^Min/+^*;*Il10^-/-^* colon tumor data re-plotted from Figure 1B are shown as white points. Each point represents data from one mouse. N= 3-5 mice per group. **(E)** 16S rRNA Shannon diversity of the IBD2, HD2, and CRC1 microbiota sampled at various time points (weeks post-gavage). N=3-5 mice per group. **(F)** Relative abundances of 11 genera by metagenomic profiling in mice colonized with the HD3, IBD3, and IBD4 microbiota across various time points (weeks post-gavage). The bars are colored according to genus name. N=3 mice per group. Statistical significance was calculated using Kruskal–Wallis followed by Mann–Whitney tests (**p< 0.01, * p< 0.05).

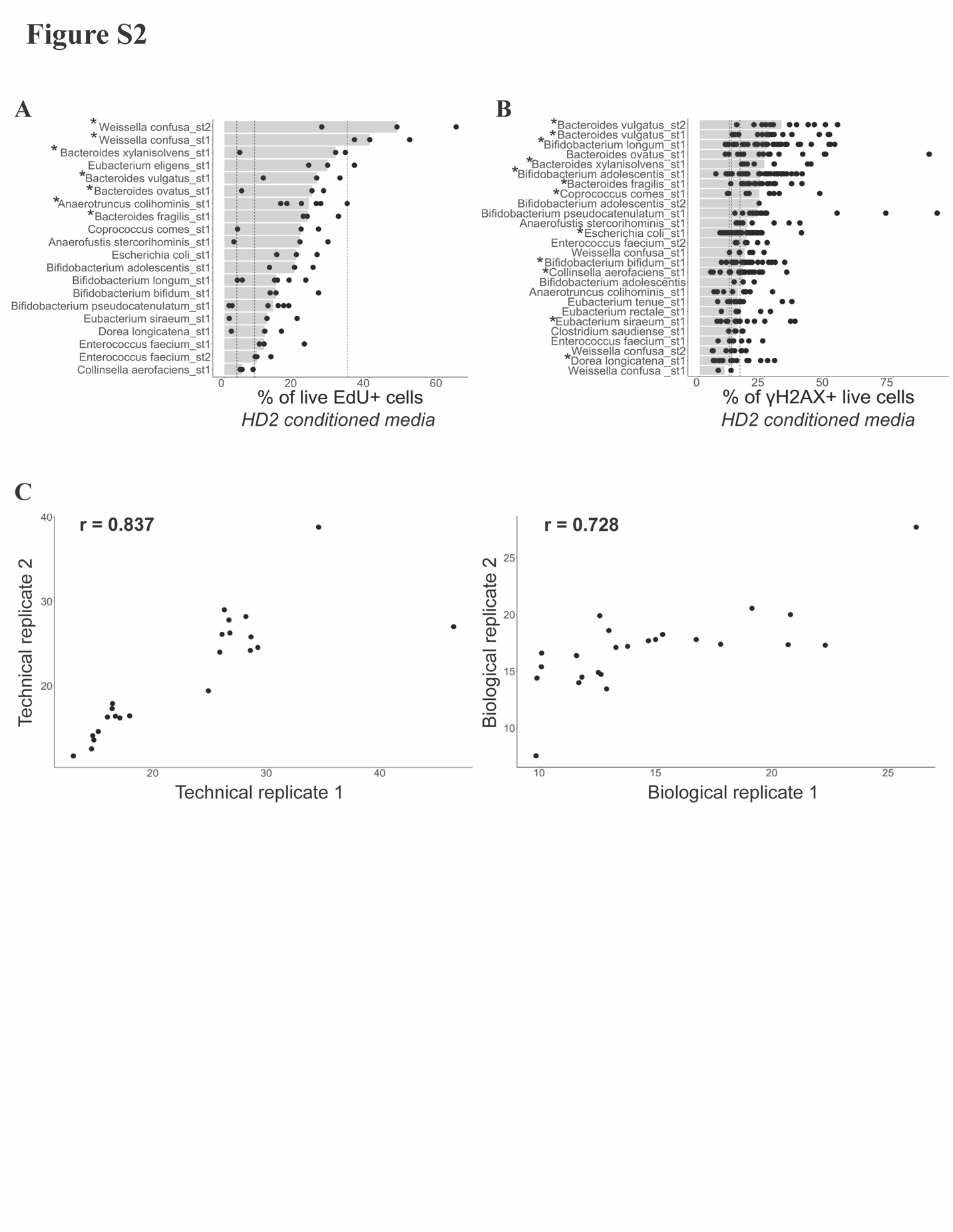

**Fig S2:** **(A)** Proportion (% of live) of EdU staining of HT-29 cells co-cultured with the stationary phase conditioned media of the HD2 bacteria cultures (presented in rank order). Each bar represents the median proportion of EdU incorporation induced by a particular condition*.* Dashed vertical line represents the proportion of EdU+ HT-29 cells treated with 1000nM ST (3.3%), 500nM ST (8.3%), or 500ng/ml EGF (33.8%) (from left to right). **(B)** Proportion (% of live cells) of γH2AX in HT-29 cells co-cultured with the stationary phase conditioned media of the HD2 bacteria cultures (presented in rank order). Each bar represents the median proportion of γH2AX induced by a particular condition*.* Dashed vertical lines represent the mean γH2AX elicited by the live *E. coli* NC101 (11.4%), *E. coli* MG1655 (12.3%), and ETBF (15.6%) bacteria. **(C)** Scatterplot of technical (left plot) and biological (right plot) replicates across the live bacteria and the stationary phase conditioned media screens. Correlation coefficient (r) calculated by Pearson.

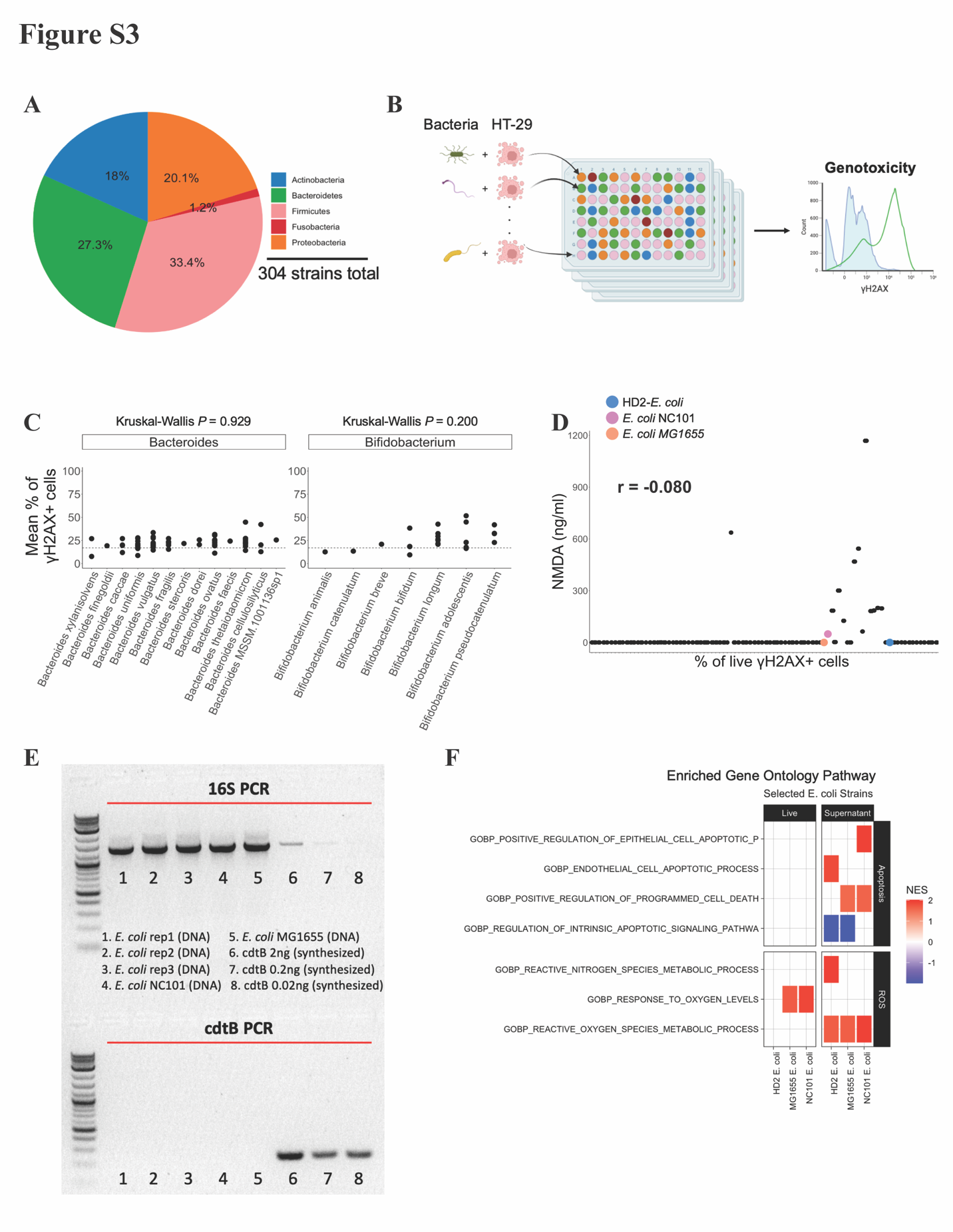

**Fig S3:** Overview of in vitro genotoxic bacteria screening methodology. **(A)** A total of 304 bacterial strains spanning the five major phyla of the human gut microbiota were cultured from 7 HDs, 5 IBD donors, and 5 CRC donors. Additional *M. morganii*, *B. fragilis*, and *E. coli* strains were included as controls. **(B)** HT-29 cells were co-cultured with unique gut-derived bacterial strains or the corresponding stationary phase conditioned media. After 24 hours, HT-29 cells were processed for intracellular γH2AX via flow cytometry. Figure created with BioRender.com. **(C)** Mean proportion (% of live cells) of γH2AX in HT-29 cells co-cultured with the live bacteria, or the stationary phase conditioned media from the 304-strain master library cultures across the two of the four most common genera. Black dashed horizontal line represents the genotoxicity threshold determined by mean + 2 x SEM γH2AX induced by HT-29 cell media alone (16.9%). Each point represents the mean value from a group of 2-6 replicates. Statistical significance was calculated using Kruskal–Wallis. **(D)** The relationship between NMDA production (ng/ml) and γH2AX induction (% of live cells) by 95 bacterial strains from tumorigenic and non-tumorigenic donors. The *E. coli* strain (0 ng/ml NMDA) from the HD2 library is highlighted in blue. *E. coli* NC101 (49.08 ng/ml NMDA) and *E. coli* MG1655 (0 ng/ml NMDA) are included for reference. **(E)** PCR was used to detect *cdtB* gene in the *E. coli* strains from the HD2 library (shown are three replicates). A synthesized *cdtB* gene construct at different concentrations was used as a positive control, while *E. coli* strains NC101 and MG1655 served as negative controls. **(F)** Heat map showing gene set enrichment results for the *E. coli* bacterial strain in the HD2 library. *E. coli* strains NC101 and MG1655 served as positive and negative controls, respectively. For any given cell, the color corresponds to the Normalized Enrichment Score (NES) and indicates how strongly the corresponding bacterial strain (x axis) is predicted to belong to the DNA damage pathway (y axis) based on the pathway’s NES (red indicates a higher NES and so, more likely to reach significance, and blue indicates a lower NES and so, less likely to reach significance).

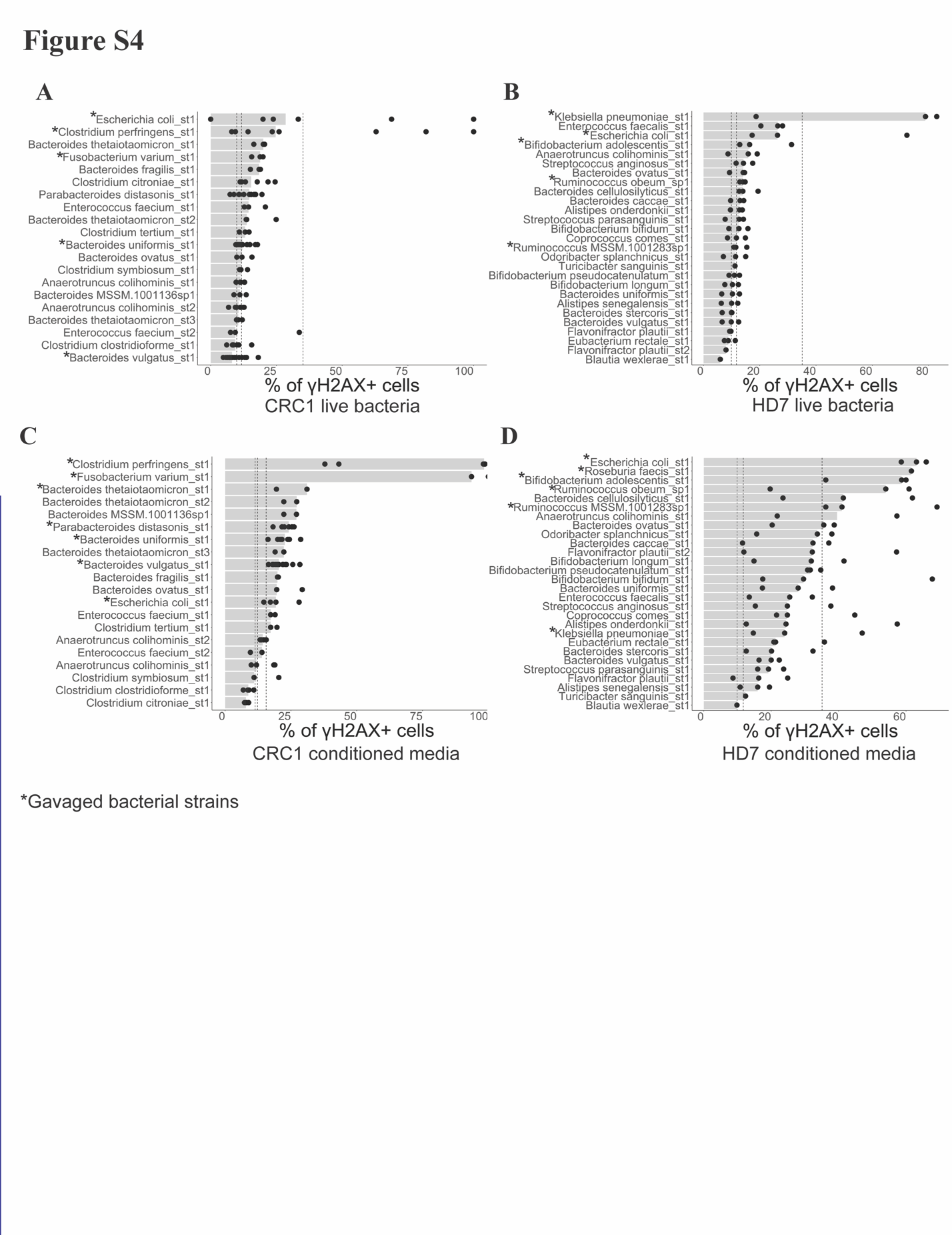

**Fig S4:** Proportion (% of live cells) of γH2AX in HT-29 cells co-cultured with the live bacteria from the **(A)** tumorigenic CRC1 or **(B)** non-tumorigenic HD7 microbiota (presented in rank order). Dashed vertical lines represent the mean γH2AX elicited by the live *E. coli* NC101 (35.1%), *E. coli* MG1655 (11.7%), and ETBF (9.9%) bacteria. Proportion (% of live cells) of γH2AX in HT-29 cells co-cultured with the stationary phase conditioned media from the **(C)** CRC1 or **(D)** HD7 microbiota (presented in rank order). Dashed vertical lines represent the mean γH2AX elicited by the stationary phase conditioned media from *E. coli* NC101 (11.4%), *E. coli* MG1655 (12.3%), and ETBF (15.6%) bacterial cultures. Asterisk next to bacteria on the y-axis indicate gavaged strains.

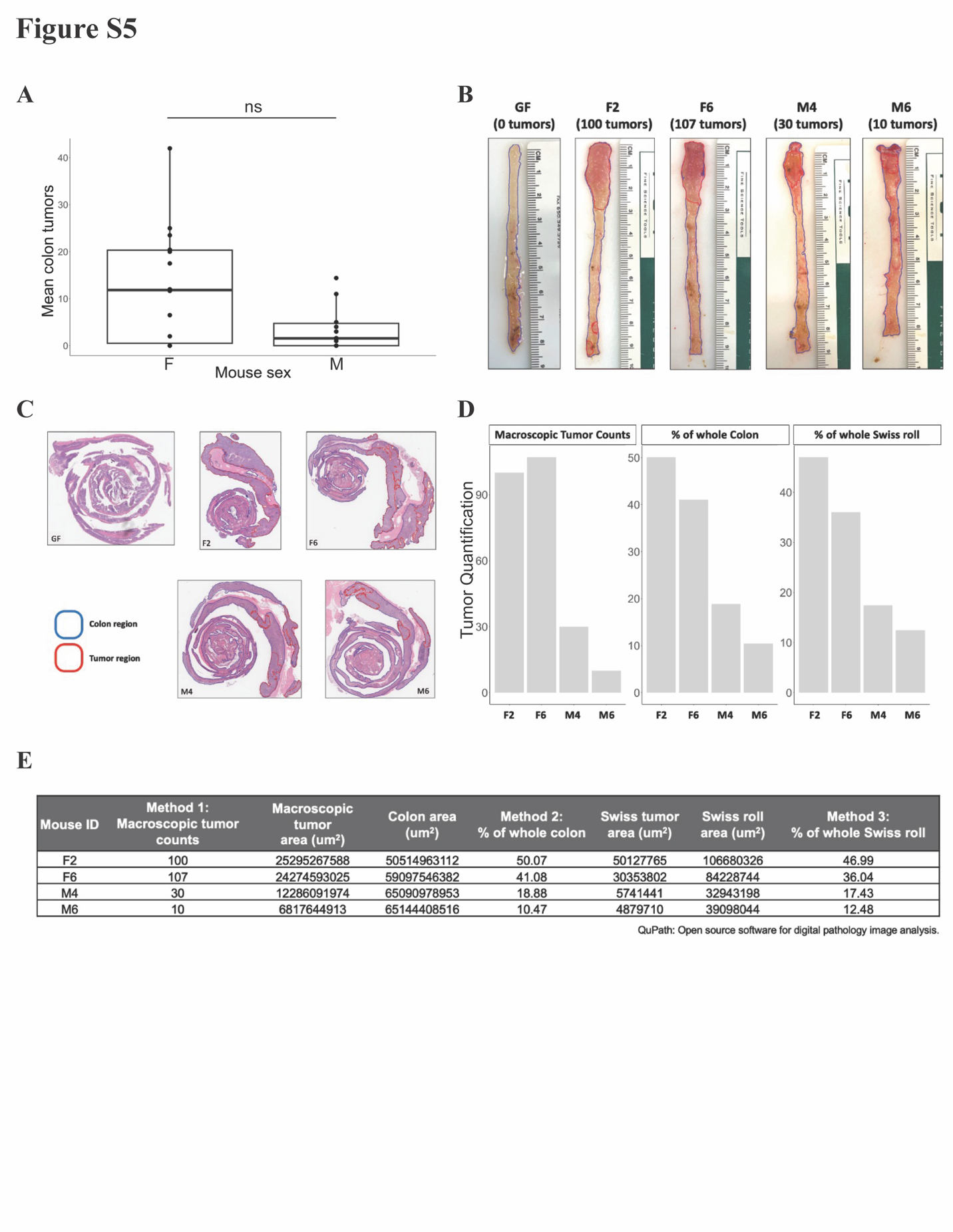

**Fig S5: (A)** Mean colon tumor counts by sex (F=Female; M=Male) of all microbiomes screened in Figure 1B. Statistical significance was calculated using Kruskal–Wallis followed by Mann–Whitney tests (ns; not significant). N=14 mice per group. **(B)** Pictures of whole colons of four mice colonized with the HD2-γH2AX+ subset, compared to a GF mouse. Mouse ID and tumor counts (shown in parenthesis) found above each picture. **(C)** Representative H&E colon sections from each of the four mice colonized with the HD2-γH2AX+ subset, compared to a GF mouse (10x magnification). Areas of tumorigenic regions and the total area of colon are outlined in red and blue, respectively. QuPath software was used to segment and measure the regions of interest. **(D)** Bar graph comparing tumor quantification using three different methodologies across mice colonized with the HD2-γH2AX+ subset. (**E)** Table detailing trends in tumor quantification across mice colonized with the HD2-γH2AX+ subset using the three different tumor counting methodologies.
